# Direct comparison of EEG resting state and task functional connectivity patterns for predicting working memory performance using connectome-based predictive modeling

**DOI:** 10.1101/2024.11.10.622847

**Authors:** Anton Pashkov, Ivan Dakhtin

## Abstract

The use of machine learning techniques combined with advanced neuroimaging methods to reveal a complex relationship between patterns of neuronal activity and behavioral characteristics is an actively developing area in modern neuroscience. Investigating brain activity data non-invasively recorded in resting-state conditions has substantially enhanced our understanding of the neuronal foundations of cognitive functions. However, results of more recent studies using functional magnetic resonance imaging suggest that task-based paradigms may outperform resting state ones in predicting cognitive outcomes. To date, no studies have tested this hypothesis using EEG data. The aim of this study was to experimentally compare, for the first time, the predictive power of models built on high-density EEG data acquired during rest and an auditory working memory task execution, while utilizing different data processing pipelines to ensure the robustness and reliability of the findings. In accordance with previous studies, we found that modeling performance was slightly better for task EEG data compared to resting state recordings, except for high working memory load, where resting state data was superior to task-based functional connectivity in terms of predictive capacity. In general, models derived from both conditions showed overall high modeling accuracy, reaching up to R^2^ = 0.31. Functional connectivity in the alpha frequency band was the most efficient predictor of working memory scores, followed by that of theta and beta bands. Finally, we demonstrated that the choice of parcellation atlas and functional connectivity method significantly impacted the ultimate results, which should be taken into consideration when designing future experiments.

## Introduction

Working memory is one of the crucial components of executive functions, allowing an organism to successfully perform a wide range of complex tasks and adapt to rapidly changing environmental demands. Over the past decade, significant progress has been made in understanding the neural mechanisms of working memory (Emch et al., 2019; Murphy et al., 2020; Wischnewski et al., 2024). An important role in this was played by the rapid development of network neuroscience and the active introduction of machine and deep learning methods into brain sciences, as well as the emergence of an independent field of research called computational neuroscience.

It is well known that any complex cognitive functions rely on the functioning of distributed neural ensembles that interact closely with each other (Christophel et al., 2017; Rezayat et al., 2022). Patterns of such activity can be non-invasively quantified by applying functional connectivity methods, which reflect the nature and degree of connection between different brain regions. However, the final outcome of such calculations directly hinges upon the choice of specific functional connectivity metrics, as well as the parcellation atlas, which determines the number of regions into which the brain is segmented (Arslan et al., 2018; Mahjoory et al., 2017).

Earlier studies have consistently and convincingly demonstrated a possibility of predicting real-life behavioral performance in a variety of cognitive tasks based on resting state neuroimaging data (Oswald et al., 2017; Rogala et al., 2020). Moving forward, more recent studies have shown a greater predictive power of task-based paradigms for forecasting specific behavioral characteristics (Finn, 2021; Zhao et al., 2023). To the best of our knowledge, all these studies have been carried out using fMRI and it remains to be determined whether these results can be applied to EEG.

In this study, we sought out to systematically compare resting state and task conditions while utilizing EEG connectome-based predictive modeling (CPM) in order to prognosticate working memory performance in a sample of young healthy subjects. In addition, we used different data processing pipelines to estimate the impact of FC method and brain atlas choice on experimental results.

## Materials and methods

### Dataset

In this study we utilized an openly available dataset (Pavlov et al., 2022), comprising resting state and task-related EEG records from 65 participants, along with demographic and behavioral data. EEG recordings were obtained using a 64-channel EEG device with active electrodes (ActiCHamp, Brain Products, Germany). The resting state data were recorded under an eyes-closed condition for a period of 4 minutes. Task-related EEG data was obtained during a working memory test involving digit spans of lengths 5, 9, and 13, presented in a random order within a single recording session. Auditory presentation of digits via speakers included a 2-second delay between digits and a 3-second delay between stimulus presentation and response. Participants were tasked with memorizing and verbally recalling as many digits as possible, resulting in a total of 108 trials. For detailed procedural information, refer to [12]. The working memory score for each participant was calculated as the number of correctly recalled digits until the first error normalized by the length of the span. Subsequently, this measure was used as a dependent variable in the connectome-based predictive modeling.

### Preprocessing

The raw data was preprocessed with MNE (Gramfort et al., 2014) and Autoreject (Jas et al., 2017) libraries. The data was filtered within a 1-45 Hz band, downsampled to 100 Hz, and rereferenced to an average electrode. Artifact rejection was fully automated, involving ICA-based ocular artifact correction and utilizing the Autoreject library to address other artifacts. Resting state data was segmented into 6-second intervals with a 1-second overlap. Task-related data was epoched into intervals of [-0.5; 2] seconds centered around the stimuli. Different span lengths (5, 9, and 13 digits) were analyzed separately, with epochs grouped based on the span length.

### Source reconstruction

The clean preprocessed data was utilized to reconstruct brain sources using the eLORETA algorithm. The fsaverage object served as the template brain model due to the unavailability of MRI scans of the participants. Two atlases were selected for brain parcellation: the Desikan-Killiany atlas (68 regions, 34 per hemisphere) and the Destrieux atlas (148 regions, 74 per hemisphere).

### Functional connectivity

Reconstructed source time-series were applied pairwise to build connectivity matrices. Six frequency bands were delineated: theta (4 - 8 Hz), low alpha (8 - 10 Hz), high alpha (10 - 13 Hz), low beta (13 - 20 Hz), high beta (20 - 30 Hz) and gamma (30 - 45 Hz). For this study, we selected 3 commonly used connectivity metrics: weighted phase lag index (wPLI, (Vinck et al., 2011)), imaginary part of coherence (imCoh, (Nolte et al., 2004)) and phase lag index (PLV, (Lachaux et al., 1999)).

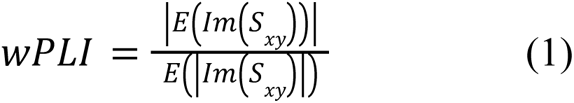

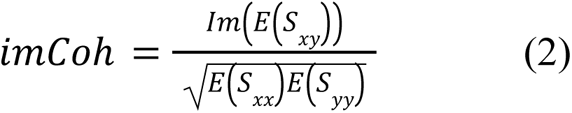

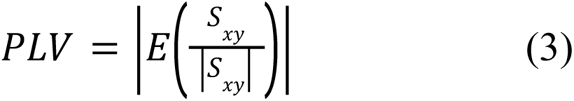

Here *S*_*xy*_ represents the cross-spectral density between signals x and y, *S*_*xx*_ (*S*_*yy*_) denotes the power spectral density of signal x (y), *E* is a mathematical expectation, *Im* is an imaginary part and |•| symbol indicates an absolute value.

### Connectome-based predictive modeling

Functional connectivity matrices were used as input for the CPM procedure (Shen et al., 2017). Given a specific edge, the procedure computes a correlation between edge weights across participants and their behavioral characteristics. If the p-value of the correlation falls behind a predetermined threshold, then the edge is marked as valuable. For each participant in the training subsample, the sum of valuable edges is calculated and these sums and the behavioral characteristics then serve as a basis to fit the regression curve. In this study, we used a threshold p-value of 0.001 and the first order of the regression curve.

### Cross-validation

The performance evaluation of each experiment was conducted using the leave-one-out cross-validation (LOOCV) technique. This method involved developing a predictive model using data from n-1 participants (the training set) to predict the score of the remaining participant (the test set) in each iteration. An experiment was considered successful if significant edges were detected in no less than 95% of LOOCV iterations. The results of successful experiments, including R-squared (R^2^, coefficient of determination) and mean absolute error (MAE), were reported. R^2^ is a quality metric that measures how well predicted values align with observed ones, compared to the «default predictor» being the mean value. Mean absolute error, on the other hand, is a metric that quantifies the deviation of the prediction from the observation. Unlike R^2^, which is a relative value, MAE is expressed in the same units as the prediction/observation.

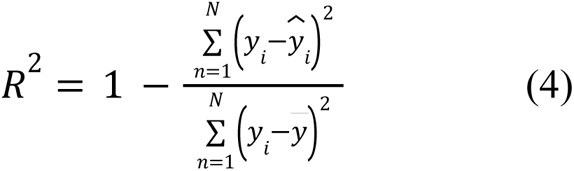

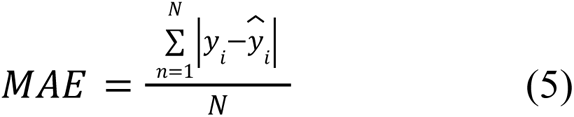

The vector *y* represents the measured working memory scores, whereas the vector 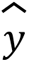 is a vector of predicted ones. 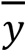 s used to denote the average working memory score, and *N* is a sample size. It is important to note that, according to this definition, R² may take on negative values (Scheinost et al., 2019).

In the context of the CPM edge selection procedure, we employed partial Pearson correlations with two covariates: sex and age. To evaluate the significance of R^2^ values, a permutation test was conducted. Specifically, we randomly shuffled the observed intelligence values 10,000 times based on the predicted intelligence values. The p-value was calculated as the proportion of permutations in which R^2^ values were equal to or greater than the original case. To address the issue of multiple comparisons, individual p-values were adjusted using the False Discovery Rate (FDR) method. Experiments that did not meet the specified threshold were excluded from further analysis.

**Figure 1.**
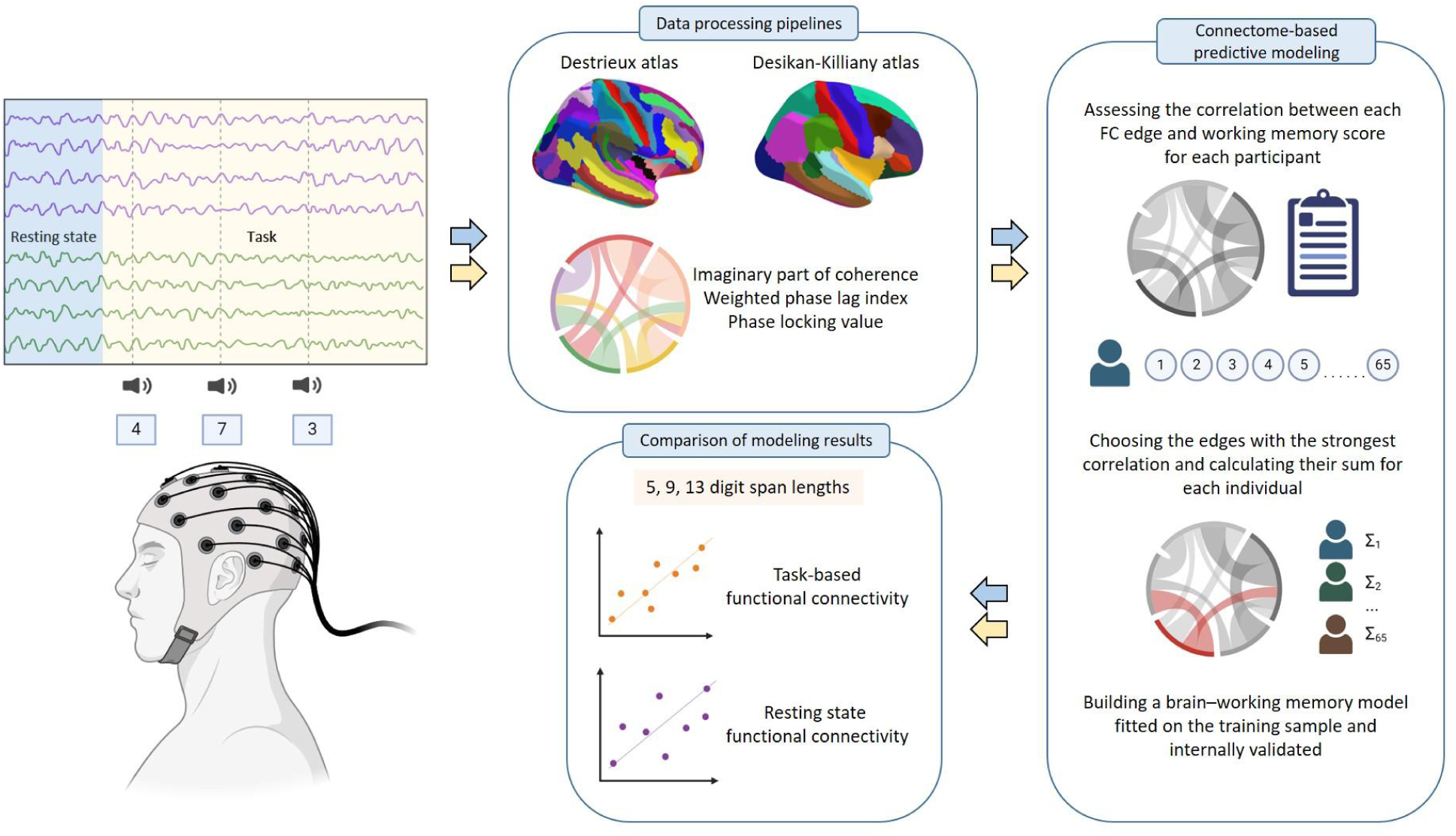
Graphical representation of the study design.

## Results

### Patients’ information and behavioral data

Continuous variables were presented as the median (interquartile range, IQR), since the data had a predominantly non-normal distribution, as determined by the Shapiro–Wilk test. A total number of participants enrolled to take part in the experiment was 86. The resting state EEG data was collected from 65 participants, while the task-related data was available only for 63 individuals (56 females, 7 males, aged 18 to 37 years, median = 20, IQR = 1). Two participants were excluded due to data processing issues. Fifty-six individuals were right-handed, six participants showed left-hand preference, and the remaining two persons were ambidextrous. Handedness was assessed using the Annett Handedness Scale.

The working memory scores for different digit span lengths were as follows: for the 5-span, the median was 0.9 with an interquartile range of 0.12; for the 9-span, the median was 0.38 (IQR = 0.17); and for the 13-span, the median was 0.19 (IQR = 0.14). These distributions are visualized in Figure 2.

**Figure 2.**
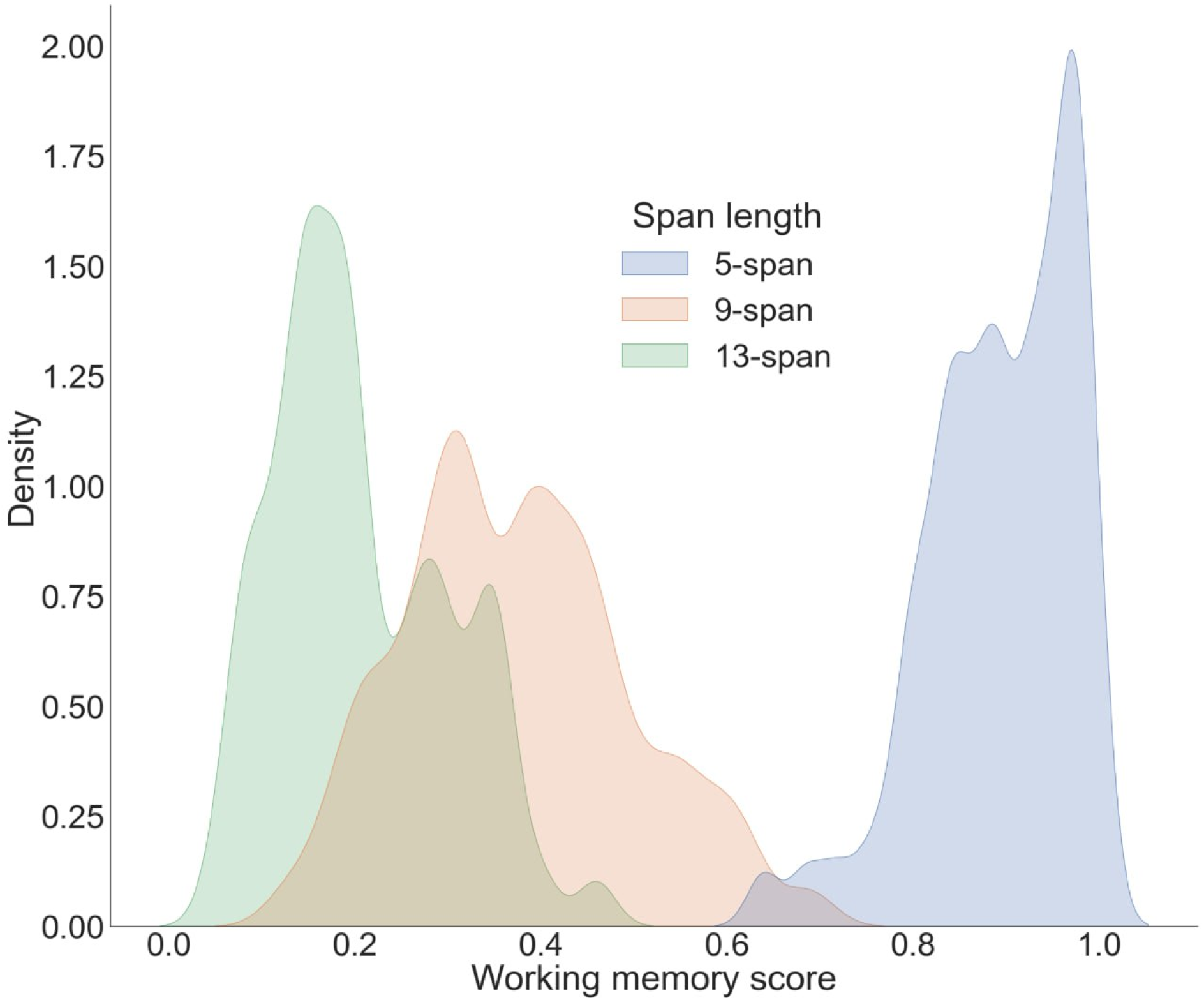
Probability density distributions of working memory scores for different digit span lengths.

### Comparison of prediction accuracy in resting state and task conditions

We conducted CPM modeling for digit sequences of varying lengths (5, 9, and 13 digits) using two different brain atlases and three distinct FC measures. The main results for 5,9 and 13 digit span conditions are summarized in Table 1. All reported p-values were FDR-adjusted.

**Table 1.** Modeling results for resting state and task-based functional connectivity EEG data. The findings are shown for two parcellation atlases and three connectivity metrics used in the study. The term «positive» refers to a positive association between functional connectivity in a specific frequency band and working memory score, while «negative» is used to indicate a negative correlation between these variables.

| Resting state connectivity | Task-based connectivity |
| --- | --- |
| 5 digit span (Desikan-Killiany atlas) |  |
| 4-8 Hz, imCoh, $R^2 = 0.11$, MAE = 0.0602 positive | 13-20 Hz, imCoh, $R^2 = 0.18$, MAE = 0.0598 negative |
| 13-20 Hz, wPLI, $R^2 = 0.07$, MAE = 0.0591 positive | |
| 5 digit span (Destrieux atlas) |  |
| No | 10-13 Hz, wPLI, $R^2 = 0.05$, MAE = 0.0603 positive |
| | 20-30 Hz, PLV, $R^2 = 0.1$, MAE = 0.0599 negative |
| | 20-30 Hz, imCoh, $R^2 = 0.1$, MAE = 0.0608 negative |
| 9 digit span (Desikan-Killiany atlas) |  |
| 4-8 Hz, wPLI, $R^2 = 0.22$, MAE = 0.0845 negative | 8-10 Hz, imCoh, $R^2 = 0.26$, MAE = 0.0842 negative |
| | 20-30 Hz, wPLI, $R^2 = 0.22$, MAE = 0.0826 positive |
| 9 digit span (Destrieux atlas) |  |
| 4-8 Hz, wPLI, $R^2 = 0.2$, MAE = 0.0898 positive | 4-8 Hz, PLV, $R^2 = 0.17$, MAE = 0.0862 positive |
| 4-8 Hz, wPLI, $R^2 = 0.11$, MAE = 0.0913 negative | 8-10 Hz, PLV, $R^2 = 0.31$, MAE = 0.0798 positive |
| 30-45 Hz, imCoh, $R^2 = 0.07$, MAE = 0.0941 negative | 8-10 Hz, imCoh, $R^2 = 0.18$, MAE = 0.0877 negative |
| | 10-13 Hz, imCoh, $R^2 = 0.26$, MAE = |
|  | 0.087 negative |
| | 10-13 Hz, PLV, $R^2 = 0.12$, MAE = 0.0907 positive |
| | 20-30 Hz, PLV, $R^2 = 0.05$, MAE = 0.0969 positive |
| 13 digit span (Desikan-Killiany atlas) |  |
| 13-20 Hz, imCoh, $R^2 = 0.13$, MAE = 0.0705 negative | 20-30 Hz, wPLI, $R^2 = 0.09$, MAE = 0.0725 positive |
| 13-20 Hz, imCoh, $R^2 = 0.12$, MAE = 0.0701 positive | |
| 13 digit span (Destrieux atlas) |  |
| 4-8 Hz, imCoh, $R^2 = 0.13$, MAE = 0.0713 negative | 20-30 Hz, imCoh, $R^2 = 0.14$, MAE = 0.0709 negative |
| 8-10 Hz, PLV, $R^2 = 0.11$, MAE = 0.0728 negative | |
| 10-13 Hz, imCoh, $R^2 = 0.09$, MAE = 0.0681 positive | |
| 10-13 Hz, imCoh, $R^2 = 0.08$, MAE = 0.069 negative | |
| 10-13 Hz, PLV, $R^2 = 0.05$, MAE = 0.0738 positive | |
| 30-45 Hz, imCoh, $R^2 = 0.28$, MAE = 0.0603 positive | |

In the resting state data obtained from the Destrieux brain atlas, we did not find any significant edges that could facilitate the prediction of task performance outcomes for a 5-digit task at a threshold no lower than R^2^ = 0.05. However, some significant results were obtained when task-based functional connectivity data was employed. Utilizing wPLI as a FC measure, we were able to attain R^2^ of 0.05 (p = 0.03) in the high alpha frequency range (10-13 Hz), whereas valuable edges in the high beta rhythm (20-30 Hz) were predictive of working memory performance at the level of R^2^ = 0.1 when using both imCoh and PLV (p = 0.01 and p = 0.013, respectively). Among the selected imCoh edges, two exhibited connections from the left central sulcus to the left supramarginal gyrus and occipital pole, while the remaining edge connected the left pericallosal sulcus to the right anterior horizontal part of the lateral fissure. The PLV-based edges were exclusively interhemispheric, featuring connections between cingular and occipital areas, as well as an additional edge linking superior temporal regions. Alternatively, using the Desikan-Killiany brain parcellation with a reduced number of regions, significant results were obtained for the theta and low beta ranges in the resting state condition (R^2^ = 0.11, p = 0.007 and R^2^ = 0.07, p = 0.008, respectively). In both cases, prediction was performed on individual edges: the left precentral to the right lateral orbitofrontal cortex for the theta band and the left temporal pole to the right superior temporal cortex for the low beta band. In addition, task-based functional connectivity allowed for a more substantial prediction of working memory scores in the 13-20 Hz range (R^2^ = 0.18, p < 0.001). We identified three valuable edges connecting the right posterior cingulate to both the right and left lateral occipital cortices as well as the right parahippocampal cortex to the ipsilateral bank of the superior temporal sulcus.

In the task condition with increased working memory demands (9 digits) resting state data analysis under Destrieux parcellation was able to reasonably prognosticate task performance based on two frequency bands: the theta range exhibited an R^2^ value of 0.2 for positive edges (p < 0.001) and R^2^ = 0.11 for negative ones (p = 0.002); suprathreshold prediction accuracy was also found in low gamma frequencies (R^2^ = 0.07, p = 0.034). A total of seven positive edges, predominantly interhemispheric, were selected in the theta band, with five of them having nodes located in the frontal lobe, specifically in the inferior frontal and orbital areas. Negative edges tended to connect cingular, temporal, and occipital regions. Furthermore, all but one gamma band connections (six out of seven) were specific to the left hemisphere and showed a tendency to cluster around parietal regions.

When CPM was performed on FC matrices obtained during task execution, a prediction accuracy was markedly improved. Specifically, the theta band positive edges demonstrated modeling accuracy of R^2^ = 0.17 (p < 0.001). These edges connected the right calcarine sulcus to the right cuneus, right suborbital sulcus to the opercular part of the left inferior frontal gyrus (IFG) and right cuneus to the dorsal posterior cingulate cortex. Moreover, both low and high alpha bands were shown to be predictive of cognitive task performance measures (imCoh 8-10 Hz: R^2^ = 0.18, p < 0.001; imCoh 10-13 Hz: R^2^ = 0.26, p < 0.001; PLV 8-10 Hz: R^2^ = 0.31, p < 0.001; PLV 10-13 Hz: R^2^ = 0.12, p = 0.002; Fig 3 and Fig 4). In the specified EEG ranges, all imCoh-based edges exhibited negative values, while PLV-based edges displayed positive values. Notably, a significant majority of PLV-based connections across the entire alpha range and imCoh-based edges in the high alpha band were predominantly localized in parieto-occipital regions and cingulo-insular cortices. Conversely, imCoh low alpha connections prominently featured frontal nodes in conjunction with temporal and occipital regions. Furthermore, a distinct set of functional connections within the high beta band demonstrated a positive correlation with working memory capacity (R^2^ = 0.05, p = 0.009). Two valuable edges were retained: right calcarine sulcus to the ipsilateral cuneus and right suborbital sulcus to the left opercular part of the inferior frontal gyrus.

**Figure 3.**
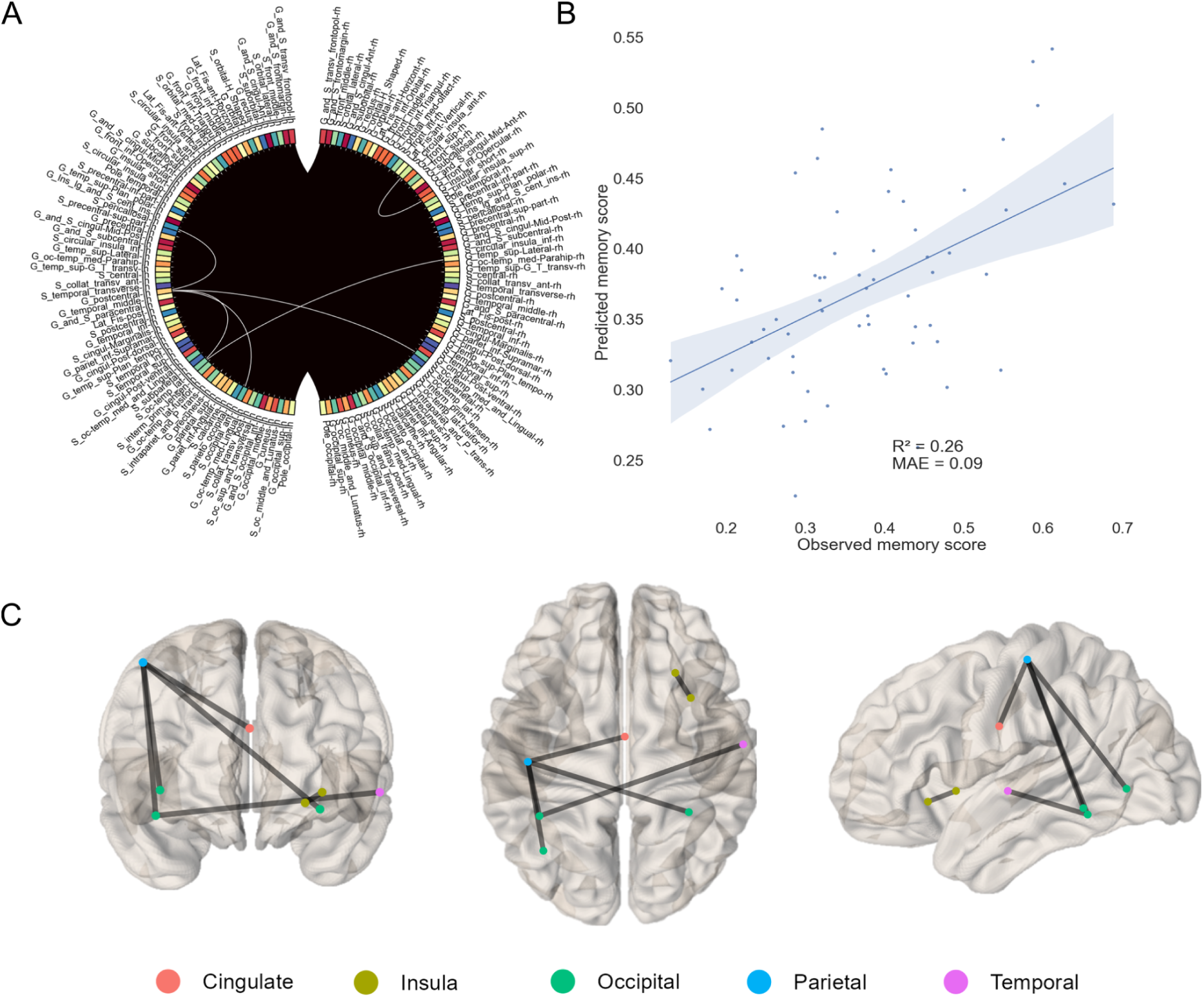
A. The circular plot representing imCoh-based valuable negative edges between brain regions parcellated according to Destrieux atlas in the high alpha (10-13 Hz) band during task performance (9-digit working memory task). B. The scatter plot displaying observed memory scores for each individual plotted against the corresponding predicted memory scores. The blue line represents the linear regression curve generated by least squares algorithm with area around the line depicting 95% confidence interval. C. Axial, horizontal and lateral views of 3D transparent brain images. The black lines display selected valuable edges and multicolored circles signify brain lobes.

**Figure 4.**
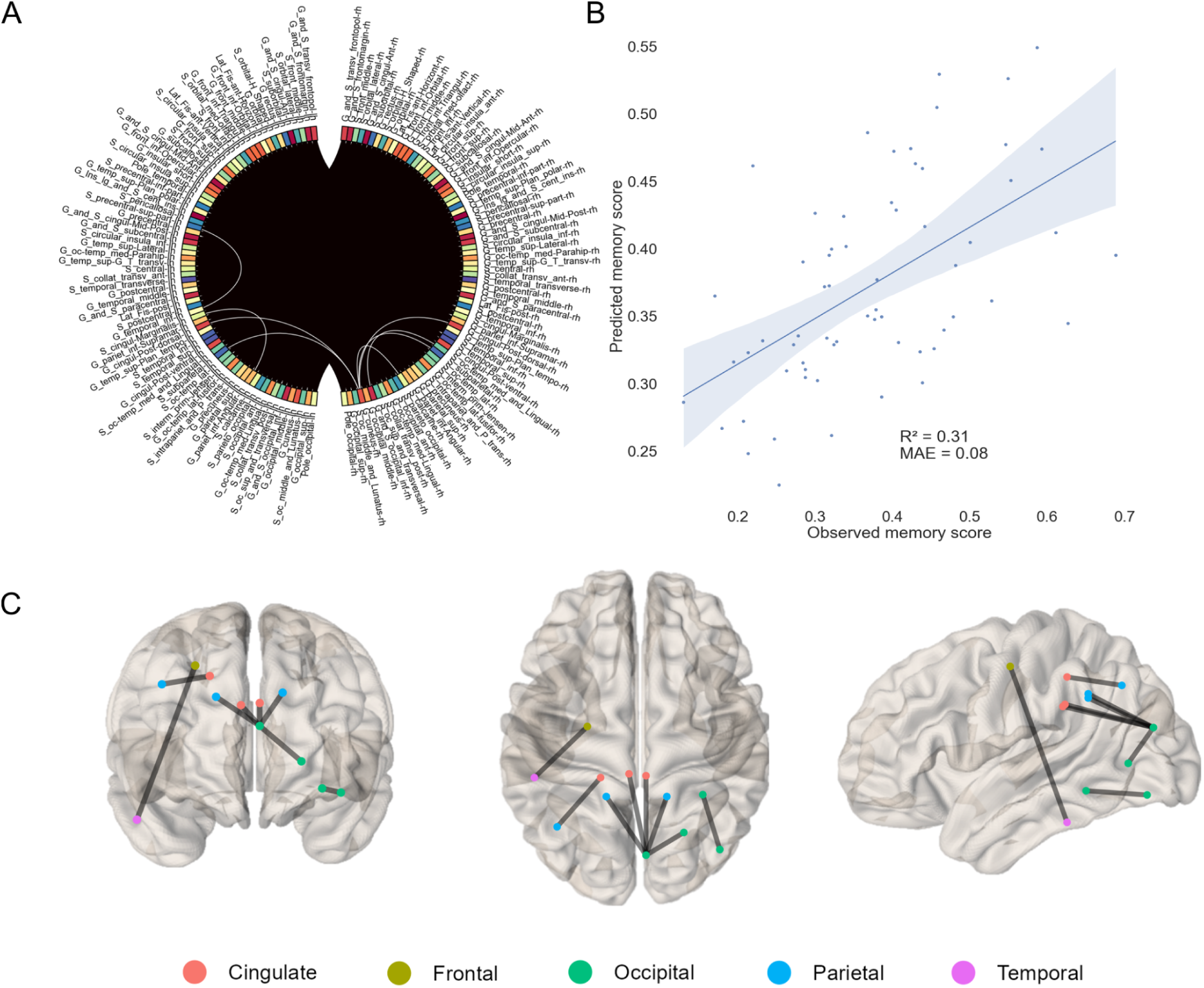
A. The circular plot represents PLV-based valuable positive edges between brain regions parcellated according to Destrieux atlas in the low alpha (8-10 Hz) band during task performance (9-digit working memory task). B. The scatter plot displaying observed memory scores for each individual plotted against the corresponding predicted memory scores. The blue line represents the linear regression curve generated by least squares algorithm with area around the line depicting 95% confidence interval. C. Axial, horizontal and lateral views of 3D transparent brain images. The black lines display selected valuable edges and multicolored circles signify brain lobes.

Switching from Destrieux to Desikan-Killiany atlas resulted in a notable decrease in the number of significant correlations between neural activity and cognitive performance. Resting-state theta band connectivity exhibited a negative correlation with working memory scores (R^2^ = 0.22, p < 0.001). Nodes in the left hemisphere were located in parieto-occipital regions, while nodes in the right hemisphere were predominantly situated in the fronto-parietal and parahippocampal cortices. Similar to findings with the Destrieux atlas, functional connectivity in the low alpha and high beta bands during task execution could forecast task outcomes (R^2^ = 0.26, p < 0.001 and R^2^ = 0.22, p < 0.001, respectively; Fig 5). Low-frequency alpha connectivity was observed between the right rostral middle frontal cortex and the pars triangularis of the IFG in the left hemisphere. Additionally, connections were found from the right IFG pars opercularis to the left insula, parahippocampal and medial orbitofrontal cortices in the right hemisphere, as well as from the right superior parietal area to the ipsilateral IFG pars triangularis. In terms of beta band connectivity, six edges were identified spanning all brain lobes except for the insular region, with only two nodes in the right hemisphere located in the transverse temporal and superior frontal cortices.

**Figure 5.**
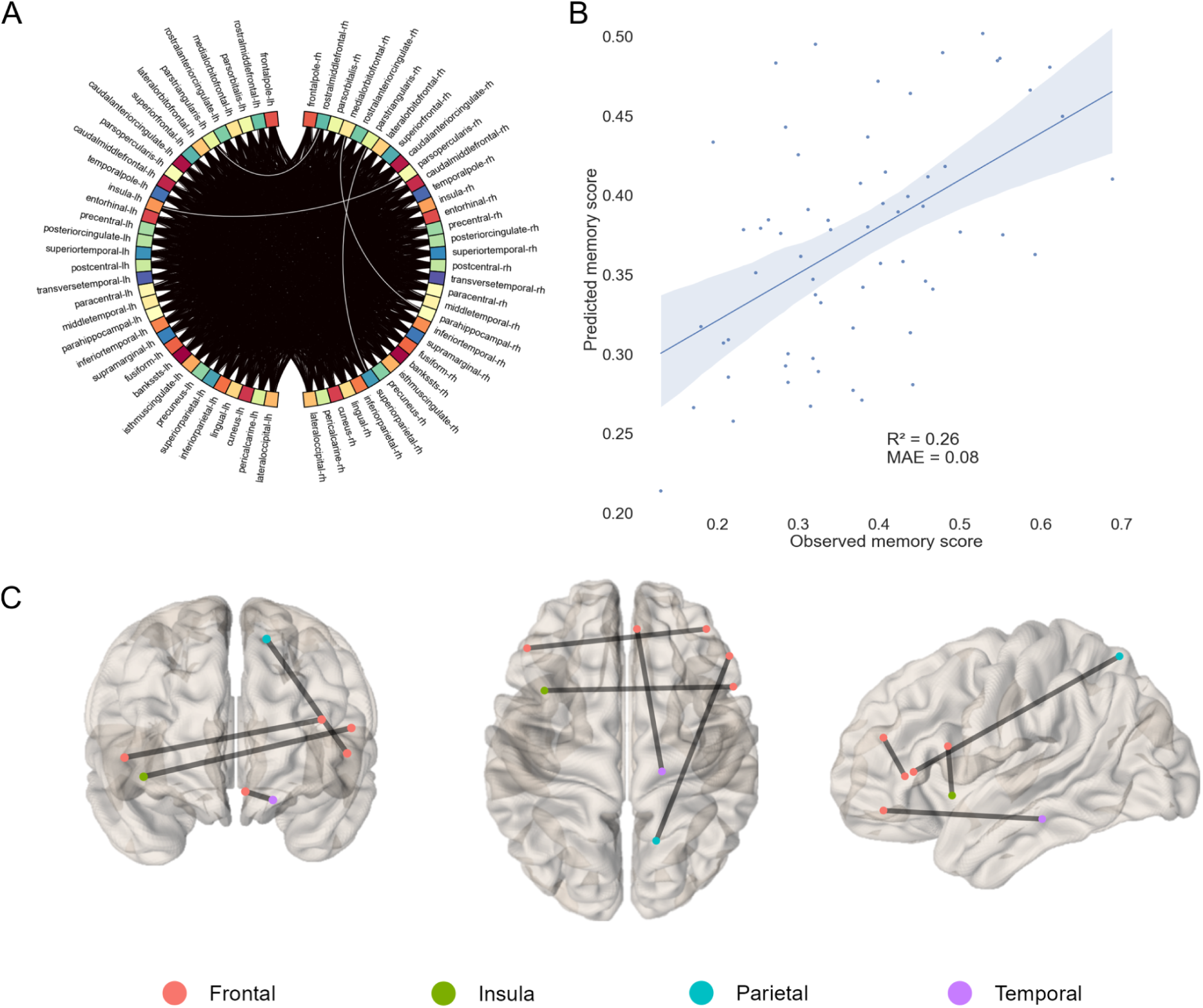
A. The circular plot representing imCoh-based valuable negative edges between brain regions parcellated according to Desikan-Killiany atlas in the low alpha (8-10 Hz) band during task performance (9-digit working memory task). B. The scatter plot displaying observed memory scores for each individual plotted against the corresponding predicted memory scores. The blue line represents the linear regression curve generated by least squares algorithm with area around the line depicting 95% confidence interval. C. Axial, horizontal and lateral views of 3D transparent brain images. The black lines display selected valuable edges and multicolored circles signify brain lobes.

Finally, a condition with the highest mental workload of 13 digits was simulated. In this scenario, resting-state EEG data using Destrieux parcellation revealed more significant and numerous brain-behavioral correlations compared to task-based recordings. Theta rhythm imCoh edges exhibited a negative correlation with working memory, yielding an R^2^ value of 0.13 (p = 0.006). Seven valuable edges were identified without any specific preference for engagement in a particular brain lobe. Low alpha band PLV edges were found to be predictive of memory scores with an R^2^ value of 0.11 (p = 0.007). Notably, this predictive capability was achieved through a single connection linking the right intraparietal area with the left middle frontal sulci. In addition, both imCoh- and PLV-based edges in the high alpha frequency range had a moderate prognostic capacity (imCoh: R^2^ = 0.09, p = 0.005 and R^2^ = 0.08, p = 0.006; PLV: R^2^ = 0.05, p = 0.025). These edges were predominantly intrahemispheric, with seven positive and five negative imCoh-based connections identified. Furthermore, two PLV-based high alpha edges were identified as contributing to the obtained R^2^ value: one connecting the left lingual sulcus to the right transverse temporal cortex, and another linking the transverse temporal area to the inferior temporal cortex. Notably, the analysis highlighted that the most significant predictive power was associated with the 30-45 Hz frequency range (R^2^ = 0.28, p < 0.001; Fig 6). Among these findings, seven positive edges were observed spanning across cingulo-insular and occipital regions. The only significant association between cognitive functioning and neural oscillatory activity recorded during task performance was detected in the high beta band (R^2^ = 0.14, p = 0.002). We were able to detect 3 significant links in the right hemisphere connecting precentral sulcus to angular gyrus, inferior orbitofrontal to anterior cingulate cortex and superior frontal to inferior parietal cortices. On the other hand, Desikan-Killiany parcellation’s main findings were confined to the beta EEG rhythm. Resting state data analysis within the 13-20 Hz range yielded a predictive accuracy of 0.12 for positive edges (p < 0.001) and 0.13 for negative edges (p = 0.001). The dominance of selected edges in the left hemisphere was evident in this case. The task connectivity data analysis revealed a less pronounced relationship between variables in the high beta band, with an R^2^ value of 0.09 (p = 0.009). Among the two positive connections that were retained, the right IFG pars opercularis was linked to the insular cortex on the same side, while the left precuneus exhibited a connection with the right inferior parietal cortex.

**Figure 6.**
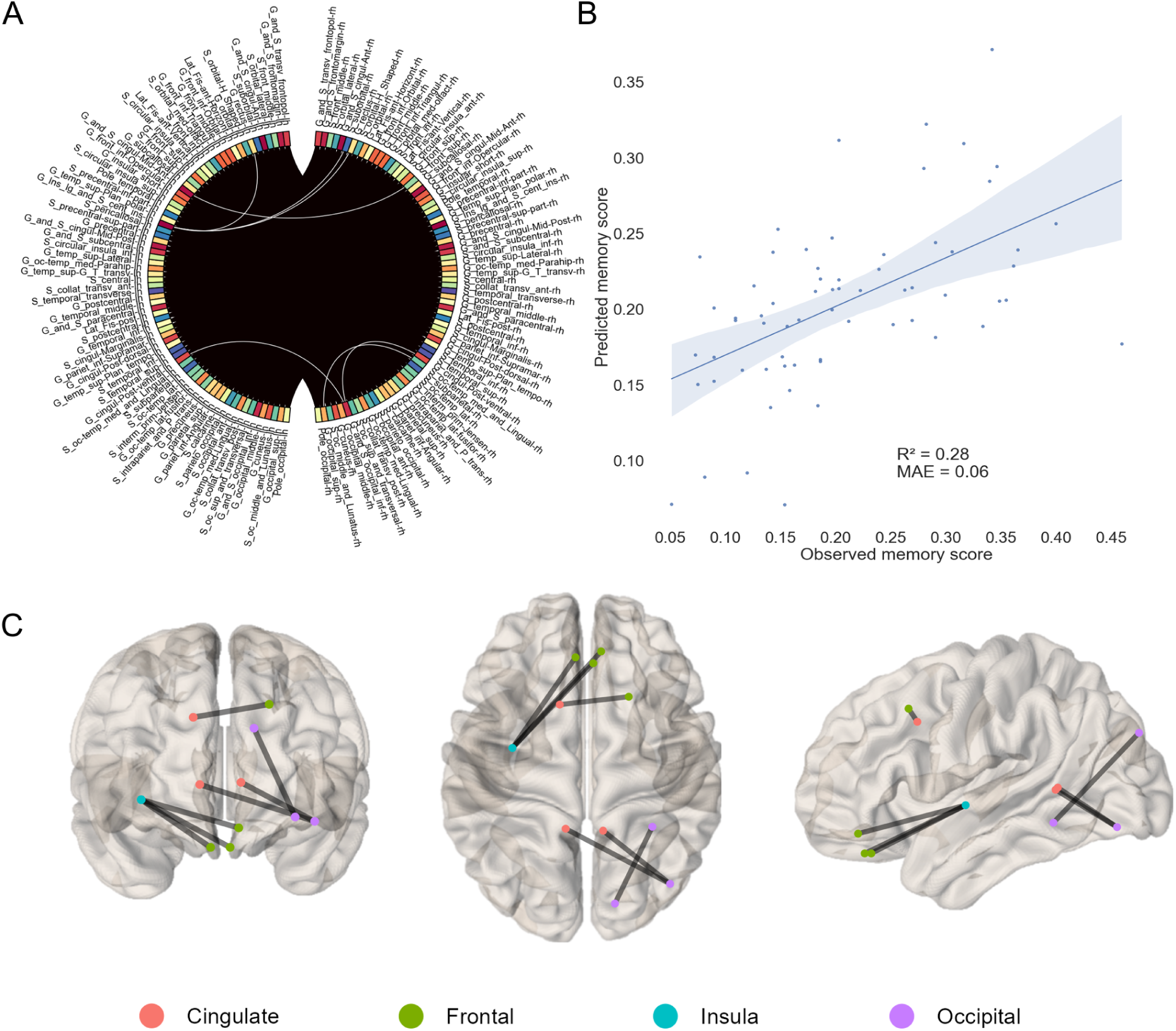
A. The circular plot representing resting state imCoh-based valuable positive edges between brain regions parcellated according to Destrieux atlas in the gamma (30-45 Hz) band (13-digit working memory task). B. The scatter plot displaying observed memory scores for each individual plotted against the corresponding predicted memory scores. The blue line represents the linear regression curve generated by least squares algorithm with area around the line depicting 95% confidence interval. C. Axial, horizontal and lateral views of 3D transparent brain images. The black lines display selected valuable edges and multicolored circles signify brain lobes.

## Discussion

In this study, we compared the performance of models run on resting state and task-based EEG connectome data in predicting individuals’ working memory scores. We found that prediction accuracy varied significantly depending on the task difficulty, parcellation atlas and functional connectivity measures used. The task set with 9 digits turned out to be the most predictive in terms of cognitive performance both in number of significant results and magnitude of prediction. Across different task and resting state settings we observed that the alpha frequency band consistently demonstrated better modeling results obtained under Destrieux parcellation, leading to a more precise prediction of working memory load. These results are in line with the available literature findings showing that alpha oscillations can be an integral part of a control mechanism over the sensory content of working memory representations (Bonnefond & Jensen, 2012; de Vries et al., 2020; Zanto et al., 2011). Moreover, it has also been demonstrated that alpha band connectivity between frontal and parietal brain areas may hold a key to adaptive working memory control (X. Chen et al., 2023). In general, brain alpha rhythm is commonly believed to implement inhibitory control over neural activity, shaping complex interactions between distant neural populations and giving rise to sophisticated intra- and internetwork balanced functioning (Jensen & Mazaheri, 2010; Klimesch, 2012). Notably, low (8-10 Hz) and high (10-13 Hz) alpha frequencies in our dataset differed in resultant modeling accuracy. Prognostic capacity of the low alpha range was overall higher than that of the high alpha. Earlier studies proposed that lower alpha range may be more concerned with attentive processes, whereas upper alpha frequencies tie more directly to mnestic functions (Klimesch, 1997).

On the other hand, application of Desikan-Killiany atlas resulted in a somewhat different set of findings. Specifically, we discovered that beta band connectivity was highly predictive of working memory load. Prior studies have provided valuable insights regarding the computational roles that can be ascribed to this frequency band during cognitive task execution (Bogaerts et al., 2020; Engel & Fries, 2010). Increase of beta activity in temporal regions has previously been demonstrated for auditory working memory tasks (Y. Chen & Huang, 2016; Leiberg et al., 2006). In our study, we showed an involvement of these regions as well. According to the state-of-the-art framework, detailing possible contribution of beta band computations to cognitive processes, this 13-30 Hz dynamics may reflect active maintenance of working memory representations (Schmidt et al., 2019). Both low (13-20 Hz) and high (20-30 Hz) beta bands in our study could predict working memory capacity reasonably well (up to R^2^ = 0.22). But these associations were valid only for the Desikan-Killiany atlas. In a recent study, it was discovered that the power of low beta frequencies allowed researchers to differentiate between successful and failed memory trials prior to stimulus onset (Scholz et al., 2017). Additionally, it has recently been reported that temporal predictions for sensory stimuli are linked to low β activity (Betti et al., 2021). High beta band oscillations may serve similar functions as activity in this range is known to increase during cognitive tasks (Baravalle et al., 2018; Z. Chen et al., 2024; Vecchio et al., 2022).

Theta oscillations (4-8 Hz) are also prominently featured in studies of working memory, both in relation to power and connectivity alterations during gradually increasing mental workload. This frequency range has long been known as being crucial for the formation of both short- and long-term memories (Herweg et al., 2020). Previous EEG studies emphasized the critical role played by theta rhythm in cognitive control in general and in working memory processing in particular (Cavanagh & Frank, 2014; Pavlov & Kotchoubey, 2022). In this study, we were able to show that the brain connectivity in the 4-8 Hz range was predictive of working memory performance, mainly during resting state independent of the parcellation employed.

At the same time, we only found a couple of significant associations between gamma rhythm and working memory scores. Interestingly, both were identified in a resting state and when using the Destrieux atlas (9 digit span — R^2^ = 0.07, 13 digit span — R^2^ = 0.28). It is well-known that gamma oscillations reflect local computational processes in neural ensembles residing in close proximity to each other, as opposed to slower wave oscillations that ensure synchronous operation of distantly located neuronal groups (Ray & Maunsell, 2015). Gamma rhythm may be a key oscillatory mechanism for grouping individual elements maintained in working memory at the peak of slower rhythms (McLelland & VanRullen, 2016). Destrieux atlas, which parcellates the cortex into smaller areas compared to Desikan-Killiany atlas, is likely better suited for detecting such local neuronal interactions. However, this hypothesis requires further experimental confirmation.

Given that we identified a significant influence of different brain atlases on the experimental results, it is pertinent to note that the choice of brain parcellation scheme is now an active area of investigation among EEG/MEG researchers. Considerable efforts are made in an attempt to mitigate the discrepancies in results obtained under different parcellations (Arslan et al., 2018; Messé, 2020; Popovych et al., 2021; Revell et al., 2022). The majority of the studies are conducted using fMRI, whereas EEG investigations are scarce. The rare exception is a study of Farahibozorg and colleagues who introduced adaptive cortical parcellation algorithms enabling researchers to determine the optimal number of parcels by minimizing leakage between them (Farahibozorg et al., 2018). In their study, they explicitly compared two such algorithms, both of which converged to a solution with around 70 cortical areas. Since our findings were heavily dependent on the brain atlas used, and considering the points mentioned above, it is possible that the Desikan-Killiany parcellation is less susceptible to the signal leakage and thus better suited for EEG data with low spatial resolution. Thus, the selection of a particular parcellation scheme constitutes a crucial determinant that significantly influences the ultimate outcomes of the study.

It is noteworthy that the selection of functional connectivity methods also strongly impacted the experimental results. We employed 3 different connectivity metrics (imCoh, wPLI, PLV) with their own unique ways of measuring a relationship between activation patterns of brain regions (Fraschini et al., 2021; Nolte et al., 2020). The application of imCoh and PLV under Destrieux parcellation was more likely to yield significant predictive outcomes, whereas imCoh and wPLI-based associations between cognitive performance scores and EEG connectivity were stronger when using the Desikan–Killiany atlas. Currently, there is no consensus among scientists about which functional connectivity methods are the most accurate and dependable for assessing neural connectivity (Bakhshayesh et al., 2019; Marquetand et al., 2019; Yoshinaga et al., 2020). It is conceivable that each of the methods we used can gauge different and specific aspects of neural communication between brain regions, with none considered superior to the others. Previous research has uncovered a wide range of complementary mechanisms that the brain can utilize to effectively execute cognitive processes, including those required for maintaining working memory (Miller et al., 2018; Zylberberg & Strowbridge, 2017). Future studies may explore recently developed sophisticated frameworks for measuring connectivity in an attempt to mitigate uncertainty related to method selection (Wang et al., 2018).

Finally, the main focus of this article was to compare predictive accuracy of models derived from EEG resting state and cognitive task functional connectivity data using connectome-based predictive modeling framework. CPM is a machine learning method that is now widely used for neuroimaging data with the goal of predicting cognitive outcomes, as well as other relevant behavioral and clinical variables (Gao et al., 2019; Yoo et al., 2018). Earlier studies employing CPM on fMRI data proved effective in precisely forecasting working memory outcomes and uncovering the neural mechanisms involved in task performance (Avery et al., 2020; Zhu et al., 2021). Additionally, a growing body of work indicates that task neuroimaging data may outperform resting state one in predicting brain–behavior associations (Gal et al., 2022; Greene et al., 2018; Zhao et al., 2023). To the best of our knowledge, our current study is a first attempt to explicitly apply CPM to EEG resting state and working memory task data. In general, based on the findings described above, we can not conclusively state that task-based functional connectivity patterns consistently outperform resting state ones when it comes to the prognostication of cognitive outcomes. Nevertheless, there was still a tendency for slightly better overall predictions based on task FC data in conditions with 5 and 9 digits. It is conceivable that task engagement leads to less variable interindividual neural dynamics in comparison to resting state condition unrestrained by external cognitive demands, which might enable better predictive accuracy of the models (Finn, 2021). In contrast, in trials when working memory load increased up to 13 digits, we found that prediction accuracy based on resting state data was superior to that of task FC profiles (both in Destrieux and Desikan-Killiany atlases). A possible explanation for the observed result may be a phenomenon of cognitive overload, which potentially leads to a decrease in attention focusing, effort investment, and, as a consequence, participant’s disengagement from the task. This is indirectly supported by data from a recent study, where the authors employed pupillometry in addition to EEG (Kosachenko et al., 2023). The pupil diameter steadily increased with increasing cognitive load, reaching a peak at about 9 digits, followed by a marked decrease as the load increased. This phenomenon remains poorly understood in modern cognitive neuroscience and deserves closer examination.

It should additionally be noted that a wide range of personal characteristics, such as gender, age, educational background, occupational and social statuses, health-related issues, and the current functional state of the organism, can significantly affect functional connectivity profiles and ultimately moderate the relationship between neural activity and behavioral variables. Moreover, the type of task given to a participant can be considered a key factor in improving or worsening a model’s predictive ability. Systematic comparison of resting state and task-based data in terms of accuracy in predicting cognitive outcomes remains an underexplored area of research, especially in EEG studies. The preliminary results obtained in this study can be seen as an important step in that direction and require further replication on larger samples using different datasets before any broad conclusions can be drawn.

Despite the overall promising results which are consistent with the previous findings, it’s worth mentioning several limitations of the current study. First, given the lack of consensus among researchers, we used an arbitrary chosen threshold of 0.05 for reporting R^2^-values in the manuscript. However, it may be considered too low to indicate the presence of a significant and meaningful relationship between behavioral and neural variables. Second, the lack of external validation in this study may limit the generalizability of the results obtained. Unfortunately, we were unable to find a suitable dataset that would allow us to attempt reproducing the experimental results. Third, in this study the data analysis was limited to the frequency range of 1–45 Hz. Although this narrow focus on traditionally investigated EEG bands is conventional for EEG studies, it might prevent the discovery of novel mechanisms underlying optimal working memory performance in delta, high gamma or ultraslow oscillations, which may be relevant for cognitive computations, as has been previously shown in a number of studies (Carver et al., 2019; Demanuele et al., 2013; Rac-Lubashevsky & Kessler, 2018).

## Author contributions

Anton Pashkov: conceptualization; formal analysis; project administration; writing — original draft. Ivan Dakhtin: conceptualization, data curation; writing — review and editing.

## Ethics approval statement

Original study was conducted in accordance with the Declaration of Helsinki and was approved by the local ethics committee (Ural Federal University ethics committee).

